# Cannibalism as a mechanism to offset reproductive costs in three-spined sticklebacks

**DOI:** 10.64898/2026.01.02.697382

**Authors:** V.I. Abuwa, A. de Flamingh, E. Arredondo, A.M. Bell, T.A. Barbasch

## Abstract

Parents often go to great lengths to promote the survival of their offspring, yet in many species parents may abandon or cannibalize offspring under their care. Using three-spined sticklebacks, where males are the sole providers of energetically costly parental care, we investigated the factors driving patterns of cannibalism in two natural populations in the field. Cannibalism was prevalent in both populations: more than 70% of parenting males contained embryos in their stomach. Neither standard length nor body condition predicted the number of embryos eaten, suggesting that males are not cannibalizing based on energetic need alone. However, males cannibalized significantly more embryos from large compared to small broods, suggesting that greater access to embryos is driving high levels of cannibalism. Using microsatellites to determine parentage, we investigated whether males cannibalize embryos fertilized by other males (heterocannibalism) to mitigate the costs of parental care. In both populations males engaged in both filial and heterocannibalism, but the two populations significantly differed in the relative amount of filial versus heterocannibalism, suggesting that males in these two populations face different reproductive costs. Combined, results from this study help disentangle the complex factors contributing to cannibalism, providing insight into how animals balance reproductive costs and benefits.

## INTRODUCTION

Parents go to great lengths to promote the survival of their offspring, often devoting time, energy, and resources toward providing for their young while forgoing their own needs (Royle et al. 2012). Parental care is also costly in terms of reduced survival and future reproductive prospects of the parent. To mitigate these costs, parents may neglect, abandon, or even cannibalize offspring that are under their care, and young conspecifics offer a vulnerable and easily exploitable food resource (Hrdy 1979; Elgar and Crespi 1992). While infanticide and cannibalism of others’ offspring can be beneficial by reducing competition for resources with one’s own offspring (Hrdy 1979; Boyko and Marshall 2009), in many cases, parents will consume some or all of their own genetic offspring (Elgar and Crespi 1992; Bose 2022). This raises a perplexing evolutionary question: why would an individual invest in producing and caring for offspring and then consume them? While seemingly counterintuitive, filial cannibalism can serve as an adaptive strategy, allowing parents to balance the benefits of improved offspring success against the costs of providing care (reviewed in Bose 2022).

Total filial cannibalism, where a parent consumes all of its current offspring, is thought to be beneficial by allowing a parent to recoup some of their lost energy in favor of future reproductive opportunities. On the other hand, partial filial cannibalism, where a parent consumes only a portion of its brood, is particularly interesting from an adaptive perspective, as it can improve the survival of the remaining offspring, for example by reducing competition for care, as well as improve future reproductive opportunities for the parent by reducing the costs of caring for the current brood (Elgar and Crespi 1992). Parental care is an energetically demanding task and often limits a parent’s ability to forage or otherwise find food for itself (Trivers 1972; Smith and Wootton 1995a; Royle et al. 2012). Thus, one of the foremost adaptive explanations for partial filial cannibalism is that consuming offspring provides energetic benefits, allowing a parent to redirect this energy to the remaining offspring or to future broods (Rohwer 1978; Elgar and Crespi 1992; Manica 2002; Klug and Bonsall 2007; Bose 2022). Indeed, the mere fact that offspring are consumed rather than removed from the nest or abandoned suggests that filial cannibalism offers some energetic benefits (Matsumoto et al. 2018; Bose 2022). However, energetic benefits do not fully explain the prevalence of filial cannibalism, and empirical support linking energy reserves to cannibalism is often equivocal (Bose 2022). Studies, particularly in fishes, have found that the prevalence of cannibalism is often unrelated to food availability (e.g. Belles-Isles and Fitzgerald 1993; Lindström and Sargent 1997; Payne et al. 2002; Klug et al. 2006) or measures of physical condition (e.g. Hyatt and Ringler 1989; Bose et al. 2019).

To account for this unexplained variation, theory has generated additional hypotheses, helping guide our understanding of how factors such as brood size, offspring developmental stage, seasonal trends, and paternity influence an individual’s decision to cannibalize (Manica 2002; Bose 2022). However, empirical tests of model predictions typically focus on a single hypothesis and are carried out within a single population, limiting our understanding of the relative importance of these different drivers and their relationship to ecology (Bose 2022).

Moreover, few studies have utilized genetic approaches to determine the extent to which cannibalism in natural populations reflects filial cannibalism of a male’s own offspring versus heterocannibalism of another male’s offspring (DeWoody et al. 2001; Bose et al. 2019). While in the lab paternity can be controlled, parents in natural populations may be caring, knowingly or unknowingly, for offspring that are not their own (Alonzo 2010; Royle et al. 2012; Rangel et al. 2023). The relative frequency of filial versus heterocannibalism has the potential to shape reproductive strategies by parents to mitigate the costs of cannibalizing one’s own offspring.

Moreover, if some of the observed cannibalized offspring in empirical datasets are actually the result of heterocannibalism, it could confound the apparent costs of cannibalism and in turn our understanding of its adaptive value. These complexities highlight the need for comprehensive studies in natural populations to disentangle the factors driving patterns of cannibalism.

Male three-spined sticklebacks (*Gasterosteus aculeatus*) offer unique opportunities to generate and test plausible alternative hypotheses for the factors related to cannibalism in natural populations. Male sticklebacks are the sole providers of energetically costly parental care, which consists of males defending their nest and fanning embryos to oxygenate them (Van Iersel 1953; Smith and Wootton 1995b; Candolin 1999). After approximately one week, the embryos hatch, and males care for fry for another week by retrieving fry that stray too far from the nest (Wootton 1984). Parenting males typically do not forage during the two-week care period, and males lose a significant amount of mass while parenting (Smith and Wootton 1995b; Guderley and Guevara 1998). The high energetic cost of parental care and reduction in foraging suggests that parenting males may cannibalize from their broods to maintain their physical condition during the breeding season (Mehlis et al. 2009). Indeed, parenting male sticklebacks are voracious predators of both embryos and fry (Foster et al. 1988; Hyatt and Ringler 1989; FitzGerald 1991; Mehlis et al. 2009), with estimates indicating that a substantial part of their diet during the breeding season is made up of embryos (Hyatt and Ringler 1989; FitzGerald 1991).

While cannibalism likely provides energetic benefits (Foster et al. 1988; Mehlis et al. 2009), access to embryos is an important factor driving high rates of cannibalism. Hyatt and Ringler (1989) found that territorial males cannibalized substantially more than non-territorial males, providing indirect evidence that males cannibalize embryos from their own nests, and thus may be engaging in filial cannibalism. However, several lines of indirect evidence suggest that heterocannibalism, rather than filial cannibalism, may predominate. Sneak fertilizations, where a male releases sperm on another male’s nest during courtship, and egg-thievery, where a male steals fertilized eggs from another male’s nests to bring to his own, are common tactics, thus males may be frequently caring for another male’s offspring (Largiadèr et al. 2001; Rangel et al. 2023). Moreover, males are more likely to cannibalize from broods with a higher proportion of unrelated embryos, suggesting that males can detect and respond to low levels of paternity in their nests potentially using olfactory cues for recognition (Mehlis et al. 2008, 2010). However, to date, few studies have utilized genetic approaches to investigate the extent to which males in natural populations cannibalize their own versus another male’s genetic offspring.

Here, we tested a set of alternative nonexclusive hypotheses for the factors influencing cannibalism in two stickleback populations. First, we compared rates of cannibalism between territorial male sticklebacks with versus without embryos in their nest to confirm previous studies suggesting that parenting males have greater access to embryos and thus cannibalize more (Hyatt and Ringler 1989). Second, we tested the hypothesis that males cannibalize embryos to gain energetic benefits, allowing them to provide more care to the current brood. We predicted that large males and males in poorer condition would cannibalize more due to their greater energetic needs. Third, we tested the hypothesis that males offset some of the costs of filial cannibalism by selectively cannibalizing embryos of lower reproductive value. We predicted that males would consume more embryos from large compared to small broods, as the value of each individual embryo is lower when there are more of them. Additionally, we predicted that males cannibalize young compared to old offspring, as broods that are in an early stage of development have lower reproductive value compared to late-stage broods closer to hatching. Fourth, we tested the hypothesis that males engage in heterocannibalism by consuming embryos that were fertilized by other males, to mitigate some of the costs of filial cannibalism. Based on previous evidence that multiple paternity within a nest is common and that males detect paternity based on egg cues (Mehlis et al. 2010), we predicted that heterocannibalism would be prevalent. Uncovering the factors that relate to the relative frequency of filial versus heterocannibalism will provide insight into the costs of reproduction and open the way for future investigations into whether males are able to mitigate these costs through engaging in heterocannibalism. We tested these four hypotheses *in situ* in two natural freshwater populations in Alaska, which allowed us to assess the generality of the results and to gain insight into the potential ecological drivers of variation in cannibalism. While we did not have strong *a priori* predictions about how these two populations might differ in cannibalism rates, stickleback populations have been shown to differ in important characteristics related to reproductive decision-making, including the rates of non-paternity at the nest (Rangel et al. 2023) and the costs of parental care (Foster et al. 2008). Therefore, differences between populations, particularly in the prevalence of filial versus heterocannibalism, can test the generality of patterns and raise novel questions about the causes of variation in reproductive costs.

## METHODS

### Timeframe for detecting cannibalism

Prior to field collections, we conducted a preliminary lab study to determine the rate at which eggs and fry are digested in the stomachs of male sticklebacks. Understanding the rate of digestion was important for the field study because we needed to know how to interpret the presence of eggs or fry in the stomachs of males on the breeding grounds, i.e., the time frame in which the cannibalism occurred. The preliminary lab study revealed that embryos took longer to digest than fry and suggests that any intact embryos found within the stomachs of cannibalistic males were likely ingested that same day, and any fry detected were likely consumed within the last hour (see Appendix).

### Field measurements and collections

Sampling occurred between June 7 – June 28, 2024 in the Cook Inlet Region of Alaska, focusing on two natural populations of sticklebacks in Spirit Lake (60.593 N, 150.986 W) and Watson Lake (60.539 N, 150.467 W). Between two and six males were sampled per day, alternating between the populations every other day, resulting in a total of 40 males sampled from each population by the end of the sampling period. Each day, males with established nests were located via snorkeling and their locations marked with buoys. These males were easily identified by their bright nuptial coloration and conspicuous nesting behaviors, such as poking at the nest opening, gluing to secure and compact the nest, and aggressive behavior toward other males (Foster et al. 2008).

Once a male was located, he was gently captured with a hand net and brought to shore for phenotypic measurements. First, male standard length was measured with calipers to the nearest 0.1mm and mass to the nearest 0.01g. Next, the male was rapidly sacrificed via decapitation, and a portion of the caudal fin was clipped and preserved in 100% ethanol. The stomach cavity was dissected, and any embryos present were extracted, counted, and preserved in labeled tubes filled with 100% ethanol. Nests were collected by gently scooping them up by hand from below and placing them in a clear plastic container. On shore, nests were gently torn open with forceps and the contents examined for the presence of embryos. If embryos were present, the brood was examined with a hand lens to determine the developmental stage (Swarup 1958). If there were multiple developmental stages present within a nest, the percentage of each stage was estimated and used to calculate the average stage of the nest contents. The presence of free-swimming fry was also noted, though no fry cannibalism was detected during this study. It was difficult to accurately stage embryos dissected from the stomach due to being partially digested, but in general, embryos collected from the stomachs appeared to be fertilized and similar in stage to embryos that were in the nest. Moreover, the stomach embryos did not show signs of disease such as cloudiness or fungal growth, suggesting that males were cannibalizing viable embryos. Embryos from the nest were preserved in ethanol and counted later in the lab.

### DNA extractions

The mean number of embryos found in males’ stomachs was 4.65, with most males consuming fewer than ten embyros, but a few males (two in Spirit Lake and four in Watson Lake) had between 11 and 31 embryos in their stomachs. We chose to extract up to ten embryos from each male’s stomach as a representative sample. Our primary goal with genetic analysis was to investigate whether cannibalism represented heterocannibalism, filial cannibalism or both processes in the two populations. Our results demonstrated that genotyping this subset of embryos allowed us to detect both filial and heterocannibalism by individual males in both populations, and thus genotyping the remaining embryos would not qualitatively influence our interpretation. DNA was extracted from fin clips and stomach embryos using a modified HotSHOT protocol from Meeker et al. (2007) and Howard et al. (2023). For fin clips, samples were first cut into small pieces and placed in 100 ul of lysis buffer (25mM NaOH + 0.2mM EDTA). For embryos, samples were placed in 30 µl of lysis buffer and the chorion pierced with forceps. Next, samples were vortexed briefly and placed in a heating block at 95°C for 30 minutes, vortexing a few times throughout to facilitate lysis. The samples were then chilled on ice and one-tenth of the volume of neutralization buffer (1M Tris-HCl, pH 8.3) was added to each tube. After a quick vortex, samples were centrifuged to pellet debris and the supernatant was stored at −20°C and thawed immediately before PCR amplification.

### Microsatellite genotyping

To test whether males cannibalized offspring that were not his own, we genotyped the males’ fin clips and the embryos collected from the stomachs of cannibalistic males at nine microsatellite loci (table A1). We chose markers shown to be highly polymorphic in freshwater stickleback populations (Bay et al., 2017). Primers were labeled with three different fluorescent dyes (VIC, PET, NED) and multiplexed in 25 ul PCR reactions using the QIAGEN Multiplex PCR kit (QIAGEN Sciences, Germantown, MD, USA). Due to the relatively high concentration of fin clip DNA compared to embryo DNA, the fin clip samples were diluted four-fold prior to PCR amplification. PCR conditions followed the kit recommendations (1 cycle: 95 °C for 15 min; 35 cycles: 94°C for 30s, 56 °C for 90s, 72 °C for 60s; 1 cycle: 60 °C for 35 min) using an annealing temperature of 56°C for all primers (Bay et al., 2017). Fragment analysis was performed by the Roy J. Carver Biotechnology Center (University of Illinois Urbana Champaign).

Genemapper was used to assess allelic sizes for all individuals at each locus. Twelve polymorphic microsatellite loci (Bay et al. 2017), were initially screened in each population using the collected fin clips, but three failed to amplify in either population and were excluded. Of the remaining nine loci (table A1), we investigated whether there was evidence for null alleles (alleles that fail to amplify via PCR) based on heterozygote deficiency, scoring errors due to stutter, and large allele drop-out, using Micro-Checker software (Van Oosterhout et al. 2004).

Micro-Checker found no evidence for scoring errors or large allele dropout, however there was evidence for null alleles at some loci in each population, with four loci for Spirit Lake and six loci for Watson Lake showing no evidence for null alleles (table A1). Null alleles, even at low frequencies, can cause false exclusion of a parent when a parent-offspring pair is heterozygous for a null allele (Dakin and Avise 2004). To account for this, all fin clips underwent an additional round of amplification and genotyping to identify null alleles. When null alleles were detected, both alleles at that locus were omitted from analysis. Alleles at each locus were manually compared between each putative father and stomach embryo and paternity was estimated using a strict exclusion criterion, where for each embryo, the putative father was excluded as a genetic match if the embryo did not share at least one allele at one or more of the amplified loci.

Exclusion probabilities of the microsatellite panel were calculated for each population as the probability of excluding an unrelated individual when one parental genotype is unknown following Jamieson and Taylor (1997), using the allele frequencies based on the parental genotypes from fin clips in each population. In both populations, panels had high combined exclusion probabilities (>0.99; table A1). To test the robustness of our assignments, we performed a likelihood-based parentage analysis using COLONY 2.0 (Wang and Santure 2009). The model runs were specified to allow for polygamy in males and females and run at medium precision with unknown maternal genotypes. Males and their stomach contents were analyzed separately, such that each model run included the male genotype as the candidate father and all the embryos genotyped from his stomach as offspring. The probability of the genetic father being among the candidate males was set to 0.5, though the results were robust to parameter values of 0.25, 0.75, and 1. In general, both methods for assigning parentage were concordant, with more offspring excluded as direct descendants using COLONY 2.0 compared to matching by hand (table A2), making the strict exclusion criteria a more conservative estimate of heterocannibalism. We therefore proceeded with matching using strict exclusion, as our goal was to detect any evidence that males consumed embryos fertilized by another male.

### Statistical analysis

Analyses were performed in R version 4.2.2. For all analyses, males with any fry in their nest were excluded as we did not detect fry cannibalism in this study, potentially due to the faster digestion time of fry compared to embryos (fig. A1), and because we could not capture or accurately count the number of free-swimming fry at the nest. A small number of males without embryos in their nest contained embryos in their stomachs but these were not included in analysis as we did not have the statistical power to investigate cannibalism by non-parenting males and our primary focus was on the factors influencing cannibalism by parents.

First, to understand the factors influencing the prevalence of cannibalism in the two populations, we used Fisher’s Exact Tests to examine pairwise comparisons in cannibalism rate between the two populations and between males with and without embryos in their nest. Next, to investigate the factors that influence an individual’s decision to cannibalize in both populations, focusing on parental males that had embryos in their nest, we fit a generalized linear model with the number of embryos eaten (including 0s) as the response variable and population, body condition, standard length, brood size (number of embryos in the nest), average stage, and experimental day as predictor variables. Body condition was calculated using Fulton’s body condition index, a common physical condition indicator used in fishes, including sticklebacks, which is based on the relationship between mass and standard length (Pawelec et al. 2016).

Brood size was determined by binning the total number of embryos observed in the nest based on a natural break in the distribution of embryos (mean = 365, median = 264), such that ‘small’ broods contained less than 300 embryos and large broods contained greater than 300 embryos, which resulted in a better fit model (based on AIC) than when brood size considered a continuous variable. Experimental day was determined as the number of days since the start of collection to control for any seasonal effects on cannibalism. We did not have strong *a priori* predictions about whether there would be interactions between any of the predictors and population, so we included the interaction only if it significantly improved the fit of the model as determined with likelihood ratio tests. We used a negative binomial error distribution to account for overdispersion.

To investigate the hypothesis that larger broods reflect greater opportunity to cannibalize compared to smaller broods, and in turn whether brood size is related to male phenotypic and reproductive traits, we fit a generalized linear model with a negative binomial error distribution with total number of embryos in the nest as the response variable and population, body condition, standard length, average stage, and experimental day as predictors. Next, we investigated whether males are cannibalizing proportionally less from larger broods, as would be expected if individual embryos from larger broods are less valuable (Manica 2002). As there was evidence for overdispersion, we fit a quasibinomial model to investigate the relationship between brood size and the probability that an embryo is cannibalized.

Lastly, to compare the proportion of filial versus heterocannibalism between the two populations, we fit a binomial regression with population as the fixed effect, such that in each Bernoulli trial a “success” was a stomach embryo that resulted from filial cannibalism (shared at least one allele with the male that consumed it at all nine loci in our parentage assignment) and a “failure” was an embryo that resulted from heterocannibalism (had at least one mismatched locus where neither allele matched the male that consumed it).

## RESULTS

### Cannibalism rate was related to parenting status in both populations

Of the 40 males sampled from each population, 18 from Spirit Lake and 20 from Watson Lake (45% and 50%, respectively) had embryos in their nest. Of those parenting males, rates of cannibalism were high, with 72% of Spirit Lake males and 85% of Watson Lake males had embryos in their stomach (fig. 1A). In contrast, 16 Spirit Lake males and 6 Watson Lake males (40% and 15%) had empty nests. Rates of cannibalism by these males were low:13% and 17% of those males contained embryos in their stomach (fig. 1B), indicating that these males either had consumed their entire brood or consumed embryos by raiding another male’s nest. Cannibalistic males consumed on average, three embryos, but this ranged from a single embryo to 31 embryos consumed. There was no statistical difference in the rate of cannibalism between the two populations for males with embryos in their nest (Fisher’s Exact Test: p = 0.47) or males with empty nests (Fisher’s Exact Test: p = 0.73). However, parenting males and non-parenting males (males with empty nests) differed in their rates of cannibalism. Males with embryos in their nest had a higher incidence of cannibalism than males with empty nests (fig. 1; Fisher’s Exact Test: p < 0.0001).

**Figure 1.**
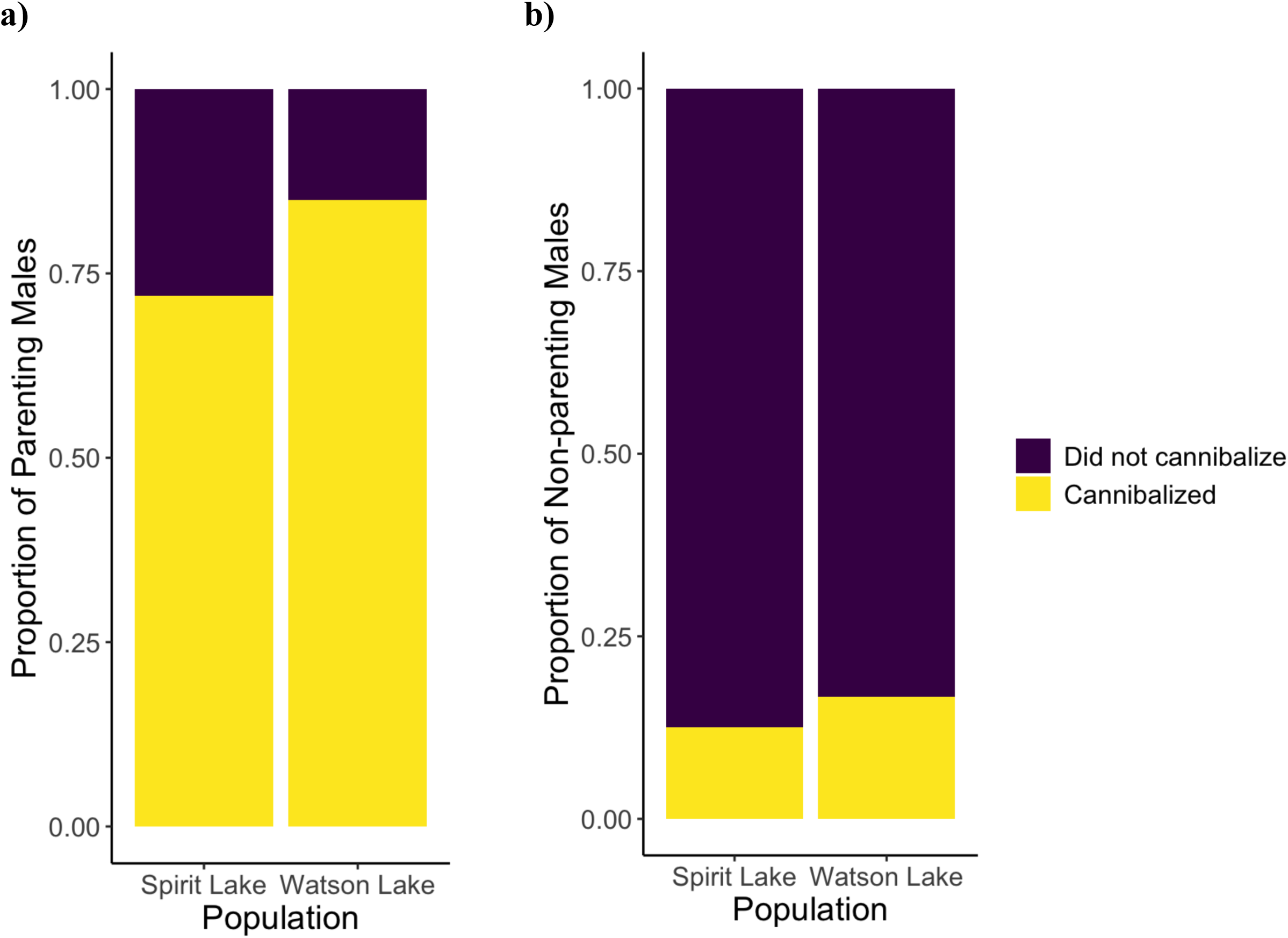
Cannibalism rates were higher in parenting compared to non-parenting males and similar in both populations. The proportion of males that had cannibalized at least one embryo for a) parenting males (n = 18 Spirit Lake, n = 20 Watson Lake) and b) non-parenting males (n = 16 Spirit Lake, n = 6 Watson Lake). Significance of comparisons between groups was determined using Fisher’s exact tests.

### Cannibalism varied among populations and was related to brood size but not energetic need

Males in the two populations did not differ in body condition, standard length, brood size, or the average stage of offspring in the nest (t-test: p-values > 0.1). Including an interaction term between population (Spirit vs Watson) and any of the other predictors did not significantly improve the fit of the model for number of embryos eaten (likelihood ratio tests: p-values > 0.1). The final model therefore included body condition, standard length, average developmental stage, brood size (small < 300; large > 300), experimental day, and population (table 1).

**Table 1.** Summary of model results for the factors associated with the number of embryos eaten and the factors associated with brood size (n = 35). Results are from negative binomial regressions including males with embryos present in their nest (i.e., parenting males).

| Model | Predictor | Estimate | se | z value | p |
| --- | --- | --- | --- | --- | --- |
| Number of embryos eaten | Fulton's body condition index | 741.97 | 617.63 | 1.20 | 0.23 |
|  | Standard Length | 0.02 | 0.05 | 0.33 | 0.74 |
|  | Developmental Stage | 0.04 | 0.03 | 1.38 | 0.17 |
|  | Brood Size (large - small) | 1.70 | 0.38 | 4.47 | **<0.0001** |
|  | Experimental Day | -0.02 | 0.03 | -0.69 | 0.49 |
|  | Population (Watson - Spirit) | 1.20 | 0.43 | 2.77 | **0.01** |
| Brood size | Fulton's body condition index | -260.98 | 482.45 | -0.54 | 0.59 |
|  | Standard Length | 0.12 | 0.03 | 3.34 | **0.0001** |
|  | Developmental Stage | -0.04 | 0.02 | -1.78 | 0.08 |
|  | Experimental Day | 0.01 | 0.03 | 0.27 | 0.78 |
|  | Population (Watson - Spirit) | -0.58 | 0.34 | -1.74 | 0.08 |

There was a significant relationship between population and the number of embryos cannibalized, indicating that males from Watson Lake cannibalized more embryos than males from Spirit Lake (fig. 2A). Contrary to our predictions, there was no relationship between the number of embryos eaten and body condition or standard length (table 1). Additionally, there was no relationship between average developmental stage and the number of embryos eaten (table 1), contrary to the prediction that males selectively cannibalize young embryos due to their low reproductive value compared to more developed embryos. However, there was a significant relationship between brood size and cannibalism, revealing that males cannibalized significantly more embryos from large compared to small broods (fig. 1B), supporting the hypothesis that males selectively cannibalize embryos of lower reproductive value, i.e., each individual embryo has lower reproductive value in larger broods. It should be noted, however, that larger broods also represent greater availability of embryos to cannibalize, and thus we can not rule out greater access to embryos as an explanation for this pattern.

**Figure 2:**
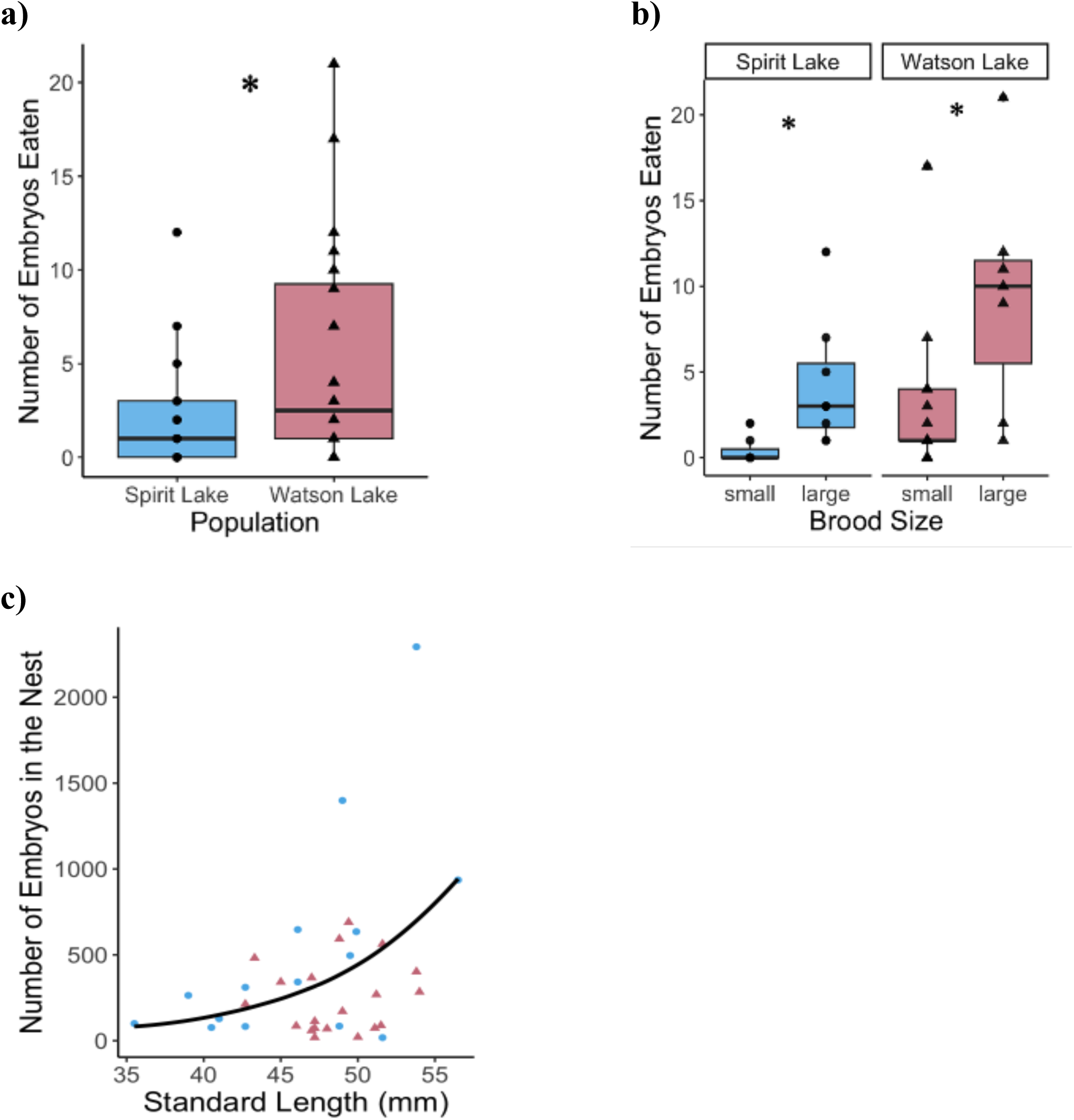
Cannibalism differed between the two populations, however within populations, cannibalism was related to greater access to embryos, with males cannibalizing more from large (>300 embryos) compared to small (<300 embryos) broods and larger males caring for larger broods. a) Males from Watson Lake consumed significantly more embryos than males from Spirit Lake. A significant positive relationship between the number of embryos eaten and a) brood size was detected. There was also a significant positive relationship between c) male standard length and brood size, but no statistically significant relationship was detected for the relationship between d) the proportion of embryos in the brood consumed and brood size. Points represent individuals, and boxplots or points are colored by population. Lines were generated using predicted values from the models for the number of embryos eaten and brood size.

### Larger males had larger broods regardless of body condition or population

Given that males cannibalize more from larger broods, we sought to identify the factors that gave males greater access to embryos and fit a model with brood size as the response variable. Including an interaction term between population and any of the predictors did not significantly improve the fit of the model for the number of embryos in the nest (likelihood ratio tests: p-values > 0.07). The final model therefore included body condition, standard length, developmental stage, experimental day, and population (table 1). The only significant effect was standard length, indicating that for every mm increase in standard length, males cared for 12% more embryos (fig. 2C). Despite the greater number of embryos eaten from large compared to small broods, a similar proportion of the brood was eaten regardless of the number of embryos in the nest (quasibinomial GLM: Estimate = −0.001, se = 0.0007, t-value = −1.62, p-value = 0.12), indicating that an individual embryo is less likely to be eaten from large broods due to the dilution effect, and thus offspring may benefit from being in larger broods.

### Heterocannibalism was common and its prevalence differed between the two populations

We used microsatellites to determine paternity in order to assess the relative contribution of filial versus heterocannibalism in the two populations. A total of 126 stomach embryos from 29 parenting males (N = 12 Spirit Lake, N = 17 Watson Lake) were genotyped. Overall, rates of heterocannibalism were high: only 8 out of 50 embryos (16%) were a genetic match to the putative father in Spirit Lake and 40 out of 86 embryos (46.5%) were a genetic match to the putative male in Watson Lake (fig. 3). Many of the embryos mismatched with the putative father at multiple loci (mismatch up to seven of the nine loci), which shows high confidence in our ability to detect heterocannibalism in our dataset (fig. 4).

**Figure 3.**
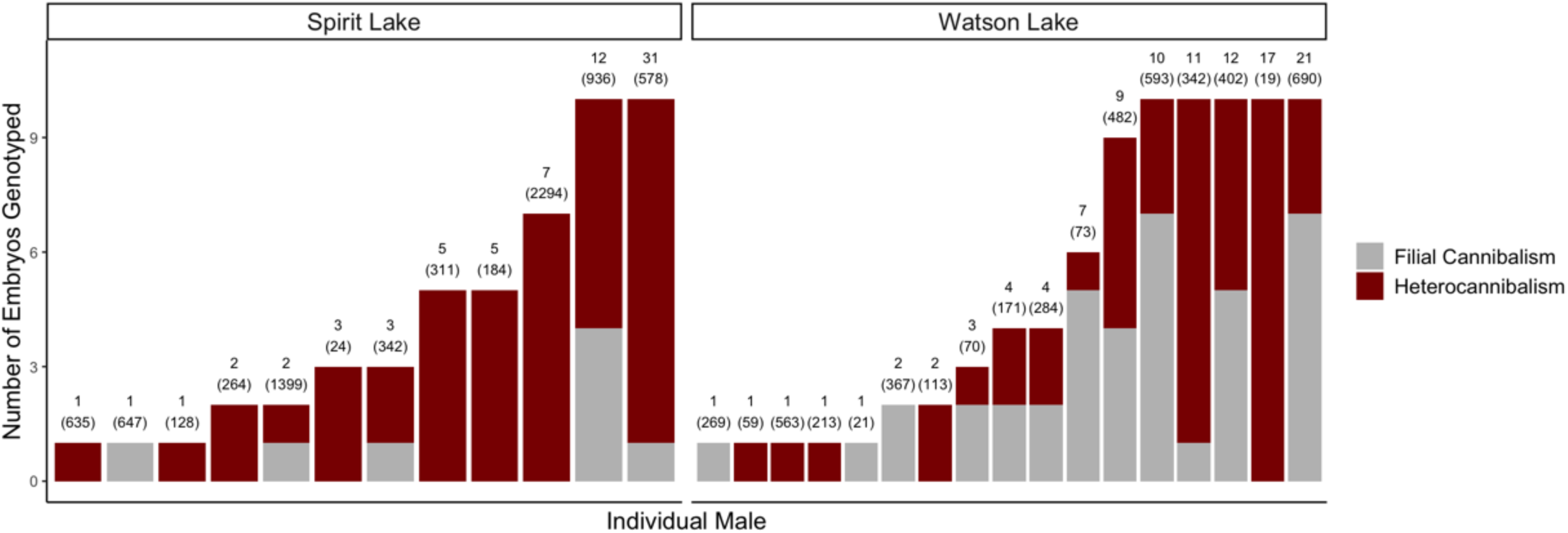
Heterocannibalism was prevalent in both populations and the populations differed in the relative amounts of filial versus heterocannibalism. Stacked bar plot showing the number of embryos eaten by individual males in Spirit Lake (left) and Watson Lake (right). Heterocannibalism (red) was determined based on a mismatch between the male and each embryo genotyped from his stomach at one or more microsatellite loci, while filial cannibalism (grey) was determined based on no mismatches at any of the nine genotyped loci. Above each bar shows the total number of embryos eaten by each male, with the total number of embryos found in his nest in parentheses.

**Figure 4.**
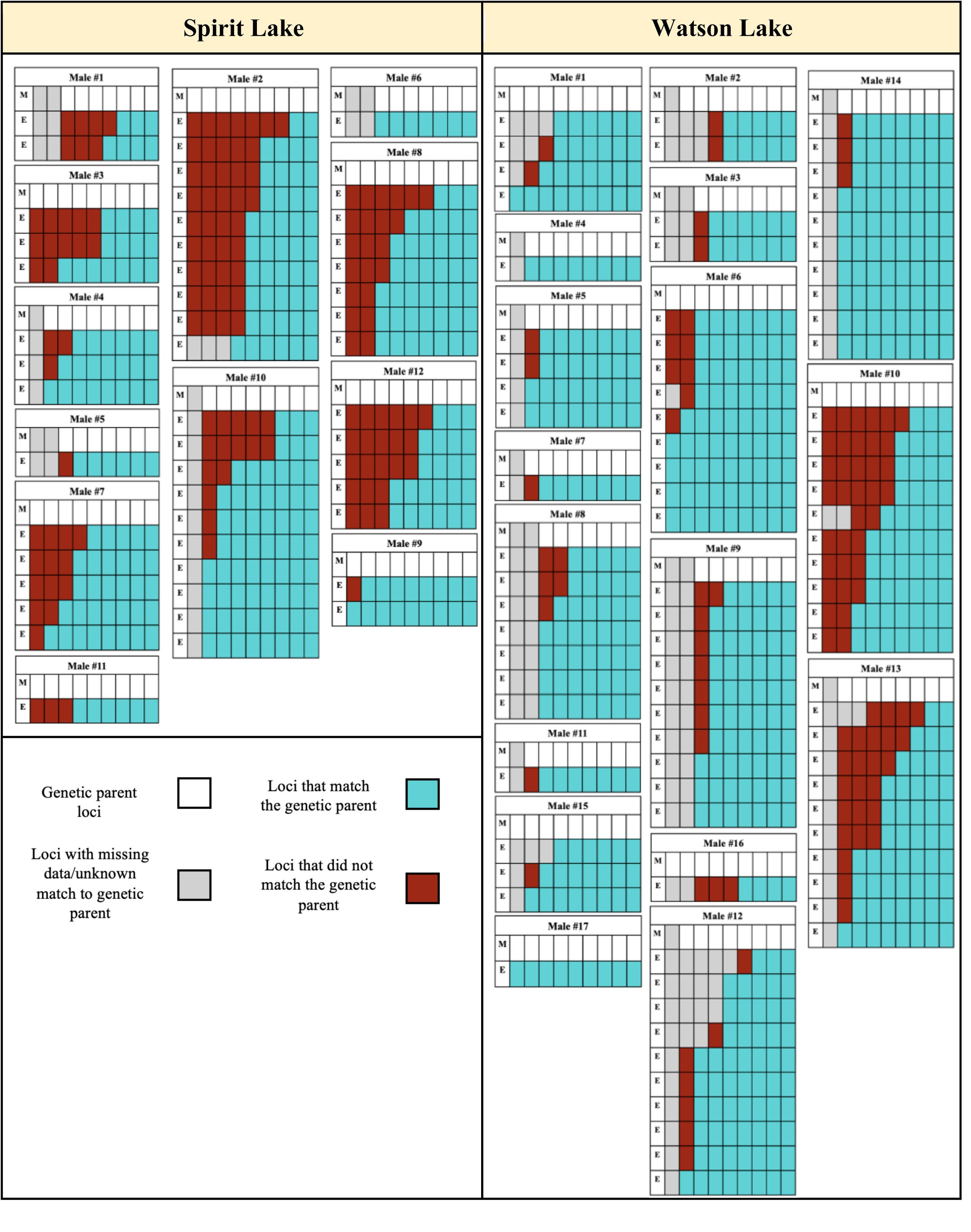
Genetic mismatch heatmap for embryos sampled from putative genetic fathers. Each box shows the genotyping results for a single male and the embryos from his stomach. The first row, labeled “M” for male, represents the loci of the putative genetic father. Each subsequent row represents one of the embryos genotyped from his stomach, labeled “E” for embryo. An individual cell represents a single locus, totaling nine in a row. White cells represent each of the father’s genotyped loci. Gray cells represent loci where a match could not be determined either because they failed to amplify from either the male or embryo sample. Blue cells indicate that the embryo’s genotype matches the father for at least one allele at that locus, while red cells mark loci where the embryo and father are genetically mismatched.

## DISCUSSION

This study investigated cannibalism in male three-spined stickleback fish to gain insight into the factors contributing to this counterintuitive behavior and more generally, into how parents balance current and future reproductive opportunities. Results showed that cannibalism of embryos by parenting males was prevalent in two natural populations of sticklebacks, consistent with other studies suggesting that embryos make up a substantial portion of sticklebacks’ diet during the breeding season (Hyatt and Ringler 1989; FitzGerald 1991; Mehlis et al. 2009). Males with embryos in their nest were more likely to cannibalize than males without embryos in their nest, and the number of embryos eaten was related to the size of the brood.

Together, these results suggest that males with greater access to embryos (caring males, males with larger broods) are more likely to engage in cannibalism. Parentage analysis further allowed us to disentangle the relative frequencies of filial and heterocannibalism and revealed that heterocannibalism was prevalent. On average, more than 85% of males had consumed at least one embryo that was not their own genetic offspring, and in many cases, the male was not the genetic father of any of the embryos he consumed, revealing that filial cannibalism cannot be inferred solely based on the presence of embryos in a parenting male’s stomach. The high prevalence of heterocannibalism is also surprising given that cannibalism by males with empty nests was rare and that cannibalism occurred even at late stages of embryonic development, suggesting that males are not only cannibalizing from their own nests, but are cannibalizing embryos they have already spent time and energy caring for. Combined, our results reveal that the fitness costs associated with cannibalism are complex and do not scale directly with the absolute number of embryos consumed, raising novel questions about the costs and benefits of parental care and cannibalism in natural populations.

Neither standard length nor body condition predicted the number of embryos eaten, which suggests that energetic need alone is not sufficient to explain cannibalism in male sticklebacks, which is one of the foremost explanations for the benefits of filial cannibalism in other taxa (Manica 2002; Bose 2022). While cannibalism likely still provides energetic benefits, the high rate of cannibalism among males in this study suggests that sticklebacks may be engaging in cannibalism to maintain their body condition throughout the breeding season, rather than cannibalizing only when in poor condition (Bose 2022). Although more embryos were eaten from large broods, the proportion of the brood that was consumed was similar across brood sizes, consistent with the hypothesis that males cannibalize more from larger compared to smaller broods because each individual embryo has lower reproductive value (Elgar and Crespi 1992; Manica 2002; Bose 2022). Additionally, we predicted that males preferentially consume young offspring due to their lower reproductive value. However, there was no relationship between developmental stage and number of embryos eaten, again supporting the hypothesis that males are consuming embryos throughout the parental care period to sustain costly caregiving behaviors, and that access to embryos, rather than energetic need or low reproductive value, is the primary driver of cannibalism. Interestingly, larger males cared for larger broods regardless of body condition or population identity. Larger males may have higher success in acquiring matings than smaller males, contributing to a larger brood size, either by mating with more females or by mating with larger females, who produce larger clutches. Indeed, both female preference for large males and male preference for large females have been demonstrated in sticklebacks (Rowland 1989; Corney and Weir 2023).

Offspring and females may also benefit from larger brood sizes due to dilution effects. From an offspring’s perspective, the likelihood of a particular individual being eaten decreases when there are more offspring in the brood. Females also potentially benefit from mating with males caring for larger broods because a female’s eggs may be less likely to be consumed (Forsgren et al. 1996). Females may gain additional benefits through mate choice for males with larger broods (or indirectly for larger males), as males with larger broods may be better able to sustain costly parental care from having greater access to embryos to consume. Given the high frequency of cannibalism among males, some of a female’s eggs could act as a kind of nuptial gift in order to compensate for the increased energetic demands associated with intense parental care behaviors, such as stronger fanning and better nest defense. Nuptial gift-giving during courtship or mating occurs in a wide variety of taxa, can take on a diversity of gift types, and can serve to balance the asymmetrical costs of reproduction between males and females (Lewis et al. 2011). In this case, laying a large clutch of eggs may be a way for the female to support the male’s parental investment while still ensuring the survival of some of her offspring.

In both populations, greater than 85% of males consumed at least one embryo fertilized by another male. Our estimates of heterocannibalism were likely conservative due to our exclusion criteria for genotyping. Heterocannibalism in sticklebacks could result from either sneak fertilizations by other males or egg stealing by the focal male. While we could not distinguish between these two mechanisms, there are several lines of evidence suggesting that males in this study may have especially cannibalized embryos that resulted from sneak fertilizations. First, males without embryos in their nest were much less likely to have cannibalized. Second, estimates from natural populations suggest that paternity at the nest is low (Rangel et al. 2023), and males are more likely to cannibalize their broods when paternity is low (Mehlis et al. 2010). Lastly, it has been suggested that when a male raids a neighboring male’s nest, he may actually be stealing eggs that he himself fertilized, as sneak spawning has been shown to precede egg stealing in the lab (Jamieson and Colgan 1992). If heterocannibalism reflects low nest paternity, our results suggest that sneak fertilizations may be incredibly common in these two populations, and that neighboring males may be caring for each other’s genetic offspring, akin to cooperative breeding systems where helpers can gain indirect or reciprocal benefits from caring for offspring that are not their own (Bergmüller et al. 2007).

While the factors related to cannibalism did not differ between the populations, males from Watson Lake consumed significantly more embryos overall compared to males from Spirit Lake. Moreover, more of the embryos eaten by males in Watson Lake were the result of filial cannibalism, suggesting that Watson Lake males face a higher reproductive cost by consuming more of their own genetic offspring compared to Spirit Lake males. It is possible that social and ecological differences between populations could help explain this pattern due to differences in the probability of sneak fertilizations and egg-thievery, both of which would increase access to embryos fertilized by other males. While not the focus of this study, the two populations studied fell along the benthic-limnetic axis of variation, which represents divergence in a suite of morphological and behavioral traits in sticklebacks that are correlated with lake size and ecology (Lavin and McPhail 1987; Foster et al. 2008; Willacker et al. 2010; Haines et al. 2023). Watson Lake is relatively shallow with murky water and has a more benthic habitat compared to Spirit Lake, which is relatively deep with clear water and more limnetic habitat (Haines et al. 2023; Hendry et al. 2024). In addition to morphological adaptations to feeding on benthic (bottom-dwelling) prey, males from benthic lakes tend to have more interactions with solitary and non-neighboring individuals and attend less to the needs of their offspring in favor of diverting intruders compared to males from limnetic lakes (Foster et al. 2008). Greater investment in nest defense could therefore increase the energetic costs of reproduction, potentially explaining the higher overall number of embryos cannibalized by the benthic Watson Lake males. Greater investment in nest defense by Watson Lake males could further serve to deter sneak fertilizations and egg-thievery, which could lead to both higher paternity at the nest and less access to embryos in other males’ nests. However, Rangel et al. (2023) found that populations with more benthic diets actually had lower paternity in their nests. Moreover, nest density was unrelated to paternity, suggesting that sneak fertilizations may not be related to the presence of nearby males. Future studies investigating the relationship between male behavior, heterocannibalism, and nest paternity would help to elucidate the mechanisms generating this variation.

An outstanding question is whether males are able to detect the paternity of offspring in their nest and in turn, whether they selectively consume embryos that were fertilized by another male. A field study of midshipman fish demonstrated that males cannibalize more from broods where paternity is low, but that males did not selectively cannibalize other males’ offspring, suggesting that males may be using indirect cues of low paternity (i.e., nest take-overs) and not direct recognition cues when making decisions about whether to cannibalize (Bose et al., 2019). In addition to indirect social cues, there is some evidence that sticklebacks may be able to use direct (phenotypic) recognition cues from eggs in their nest. Studies that eliminated social cues from rival males by manipulating paternity in the nests of parenting males found that the likelihood of total cannibalism was higher in clutches with low paternity (Frommen et al. 2007; Mehlis et al. 2010). Olfactory cues are likely to play a role in a male’s assessment of paternity (Frommen et al. 2007), and adult sticklebacks can recognize close kin using both visual and olfactory cues (Mehlis et al. 2008). It seems plausible that males cannot discriminate kinship between individual embryos, given the number of individual males in our study who had both embryos that he fertilized and embryos fertilized by another male in their stomachs. Rather, males may be using direct cues to assess overall paternity but be unable to identify specific embryos that they did not fertilize. In many mammalian taxa, paternity confusion, where the identity of the offspring’s genetic father is concealed, serves as a counterstrategy by females against infanticide by rival males (Hrdy 1979). Males in turn, may benefit from tolerating a lower probability of paternity as it reduces the risk of infanticide by other males (Boyko and Marshall 2009). Sneaking behavior in sticklebacks may serve a similar function of confusing paternity, such that parenting males must balance the risk of consuming their own offspring against the risk of caring for another male’s offspring. Alternatively, the relative frequency of filial and heterocannibalism might indicate that males resort to consuming their own genetic offspring once they consume most or all of the embryos that they did not fertilize. Future investigations comparing paternity between embryos in the stomachs and nests of cannibalistic males could help distinguish between these alternative hypotheses.

This study provides a comprehensive field-based test of alternative hypotheses to explain cannibalism in natural populations, highlighting the complexity of factors influencing reproductive costs and benefits. Rates of cannibalism were high among parenting males in our study, with the majority having cannibalized at least one embryo. This was surprising considering that our sampling only captured a snapshot in time of whether a male had cannibalized embryos that same day, suggesting that cannibalism occurs frequently and throughout the parental care period. Therefore, cannibalism may serve as a ubiquitous strategy by which males manage the costs associated with reproduction and parental care. However, our findings reveal that a high prevalence of cannibalism is not necessarily indicative of high costs associated with consuming one’s own genetic offspring. Rather, males may tolerate the costs of caring for offspring that are not their own, either because they cannot distinguish them from their own or so that he can later consume them. Additionally, differences between populations in the relative frequency of filial and heterocannibalism raise interesting questions about the social and ecological factors that shape reproductive strategies. Combined, our results help explain how parents can manage the high costs associated with reproduction and shed light on parental investment strategies and evolutionary reproductive tradeoffs.

## Notes

### Competing Interest Statement

The authors have declared no competing interest.

### Funding

- National Institutes of Health: 1R35GM139597
- U.S. National Science Foundation: 2109619

