## Supplemental text for "Cannibalism as a mechanism to offset reproductive costs in three-spined sticklebacks"

**Appendix**

*Timeframe for detecting cannibalism*

Prior to field collections, we conducted a preliminary lab study to determine the rate at which eggs and fry are digested in the stomachs of male sticklebacks (supplementary fig. A1). Understanding the rate of digestion was important for estimating when cannibalism occurred, enabling us to differentiate between same-day and consumption of embryos and fry in the past. Five males were isolated in individual tanks and starved for 24 hours before being randomly assigned to one of five experimental conditions: one, three, five, eight, or 24 hours. Using frozen, unfertilized eggs, each male was presented with five eggs and observed until all five eggs were consumed. After the allotted time per condition, males were sacrificed for stomach dissection and egg inspection. Signs of degradation and deflated chorions were recorded. The eggs in the one-hour condition sustained little to no defects and were easily countable with the human eye. After three hours, eggs exhibited signs of softening and degradation, although the chorion remained largely intact. At five hours, they were increasingly difficult to count with the naked eye and required microscopic examination. The eggs after eight hours showed signs of being digested, and no eggs remained at 24 hours. These preliminary results suggest that any intact embryos found within the stomachs of cannibalistic males were likely to have been ingested that same day.

We performed a follow-up experiment using fry based on the observation that none of the sampled males had fry in their stomach, suggesting that fry may be digested more quickly than embryos and thus more difficult to detect. Fry were collected one week after hatching and given an overdose of MS-222 and then rinsed with tank water before being presented to the focal male. Males were presented with five fry, and each individual ate between one and five fry. After one hour, fry were undetectable, thus we performed the next trials at 10, 15, 20, and 40 minutes. When only a single fry was consumed, it was digested rapidly, even after 10 minutes, and was barely detectable after 15 minutes except for the presence of undigested eyes. If multiple fry were consumed, fry were still detectable but already partially digested at 20 minutes. By 40 minutes, the fry were digested and virtually undetectable. This suggests that any fry detected in the stomachs of males were likely to have been consumed within the last hour.

|  | **10 minutes** | **15 minutes** | **20 minutes** | **40 minutes** |
| --- | --- | --- | --- | --- |
| **Fry** | 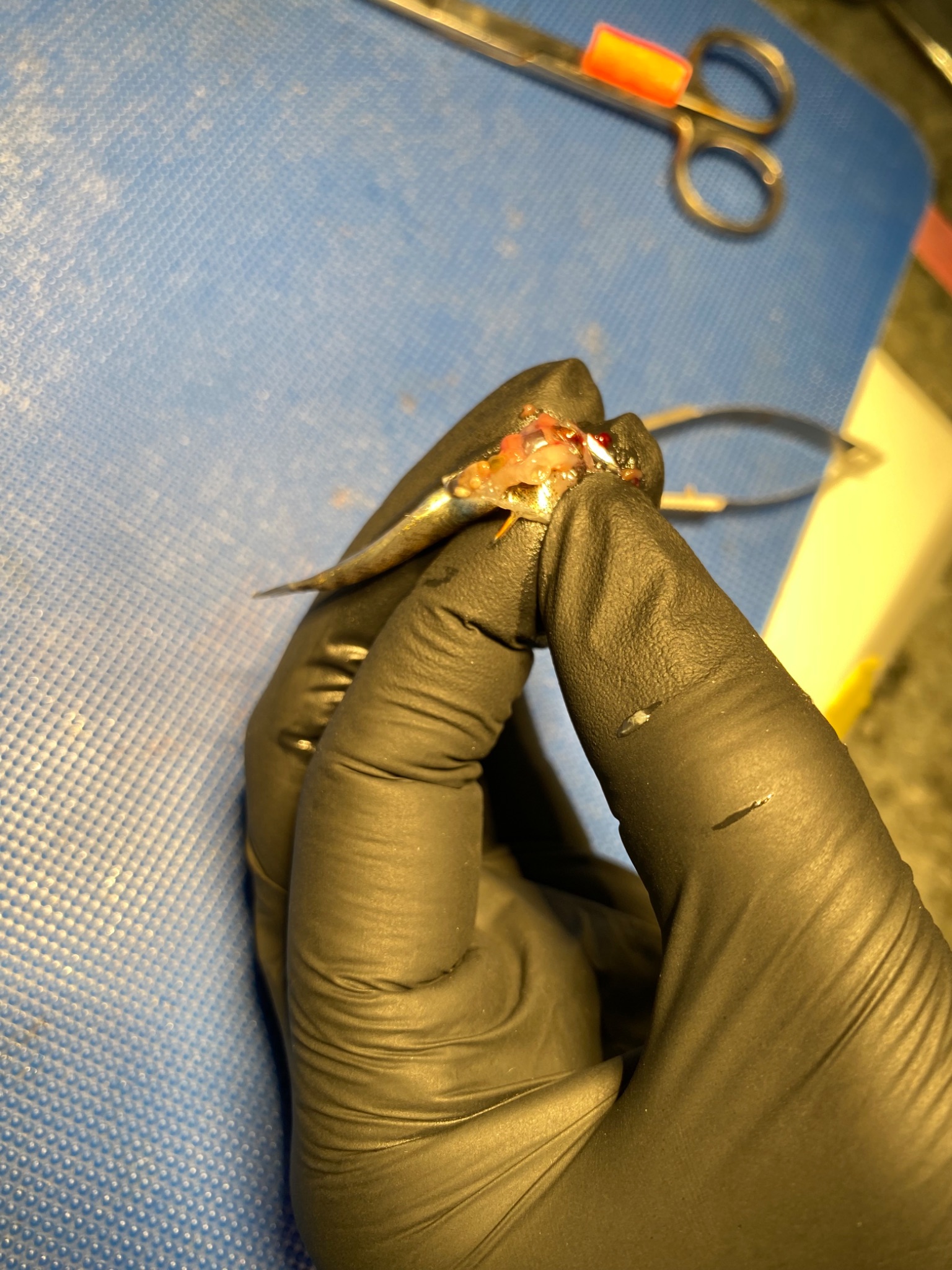 | 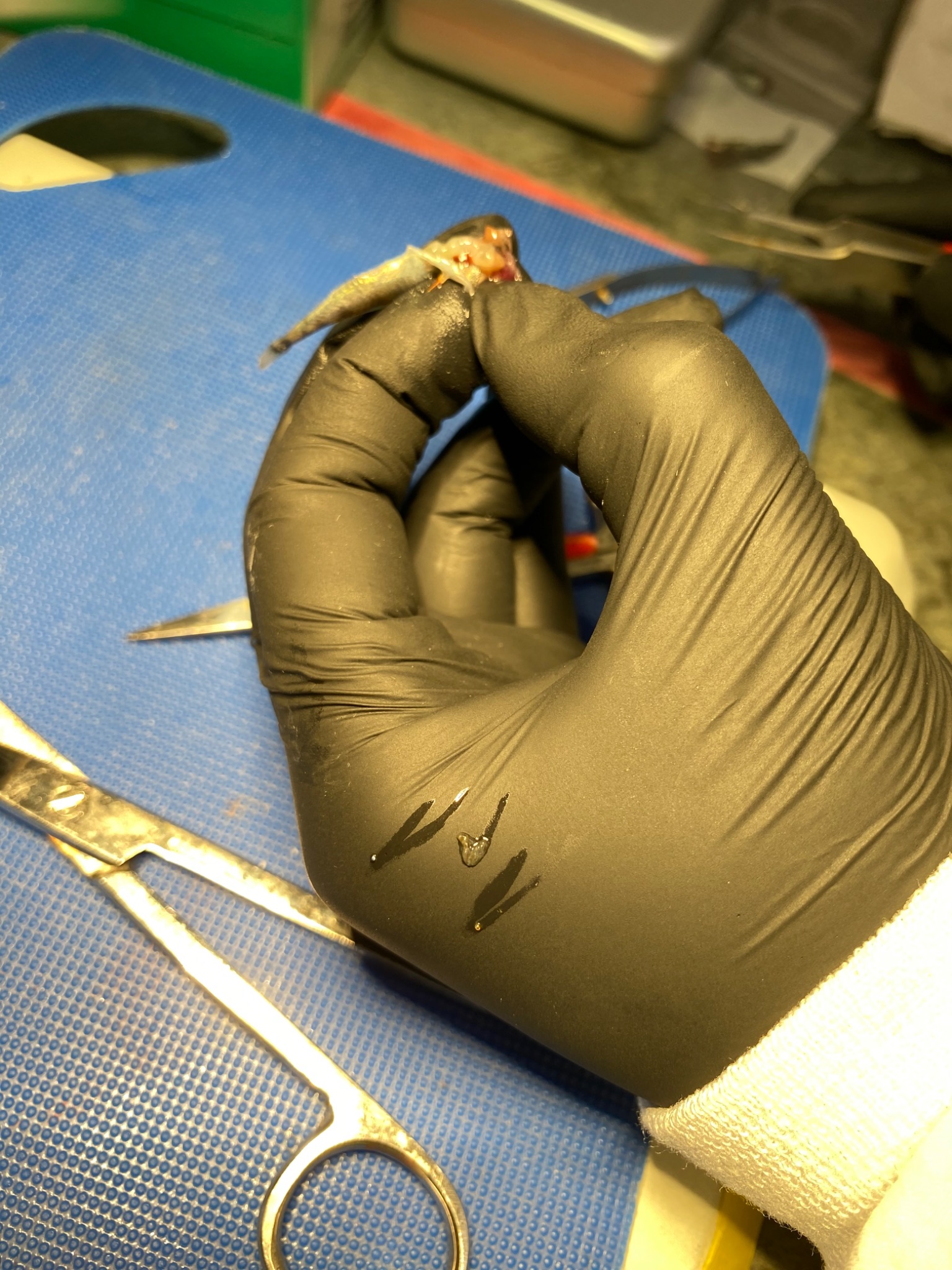 | 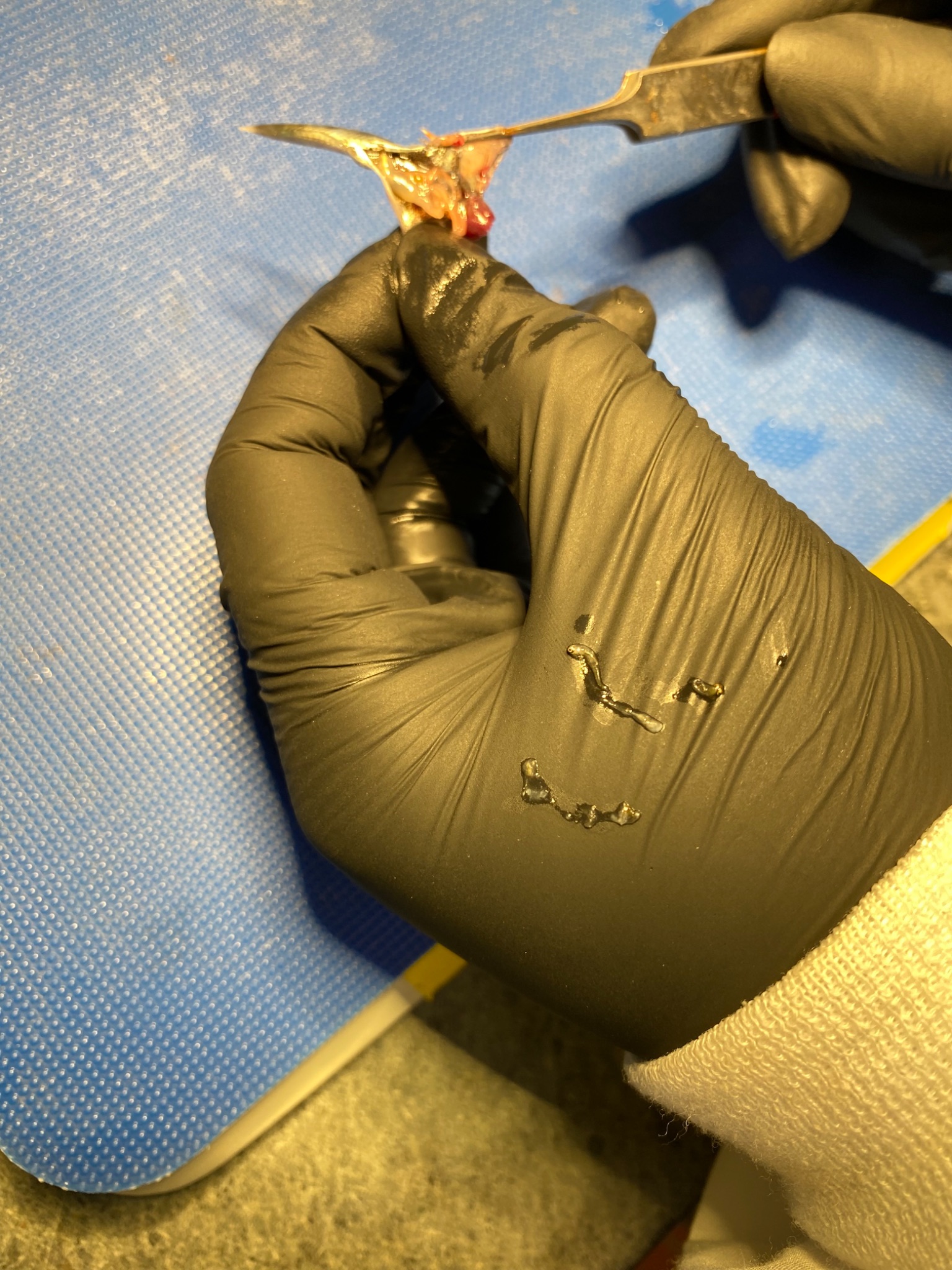 | 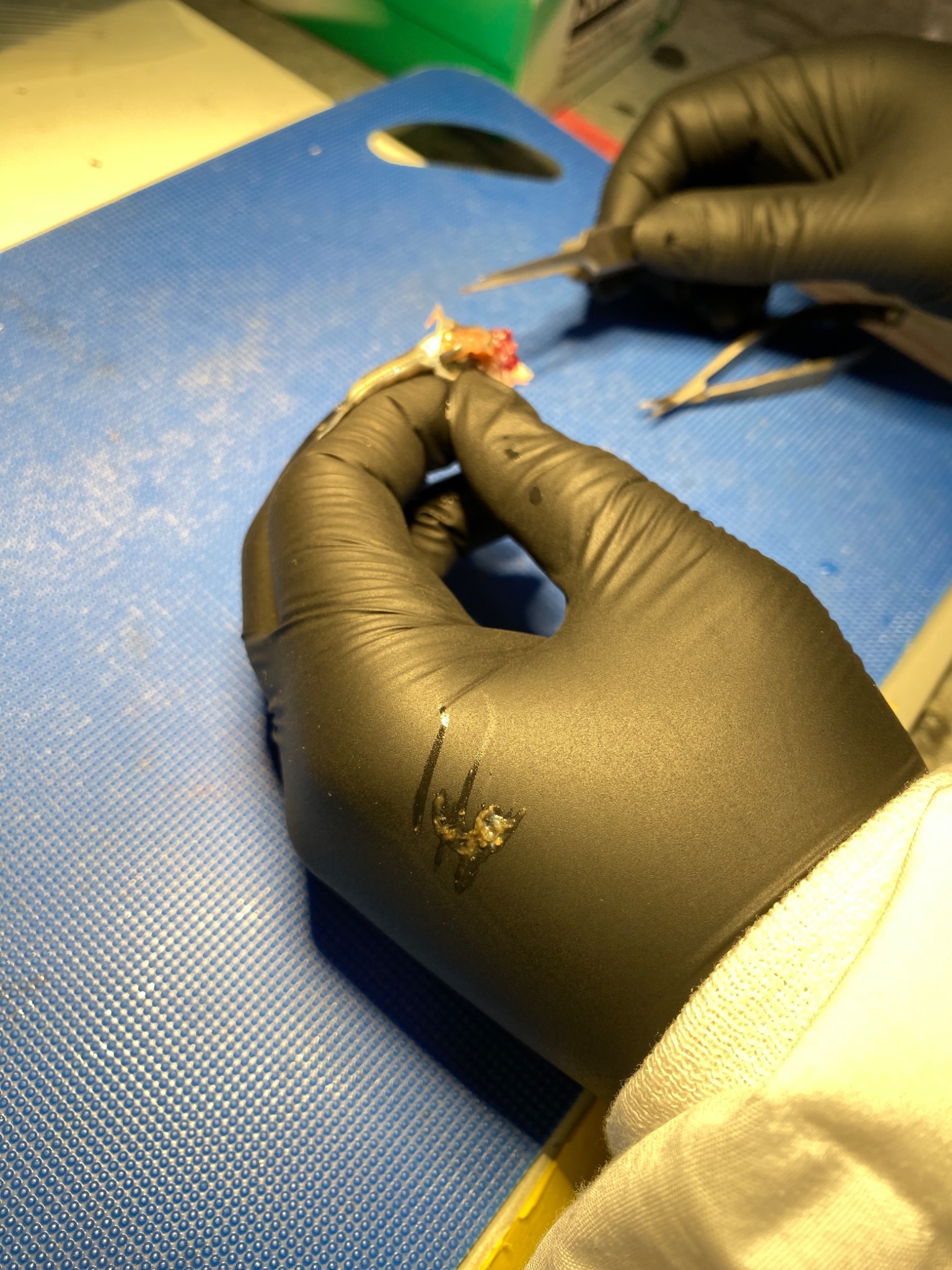 |
|  | **1 hour** | **3 hours** | **5 hours** | **8 hours** |
| **Eggs** | 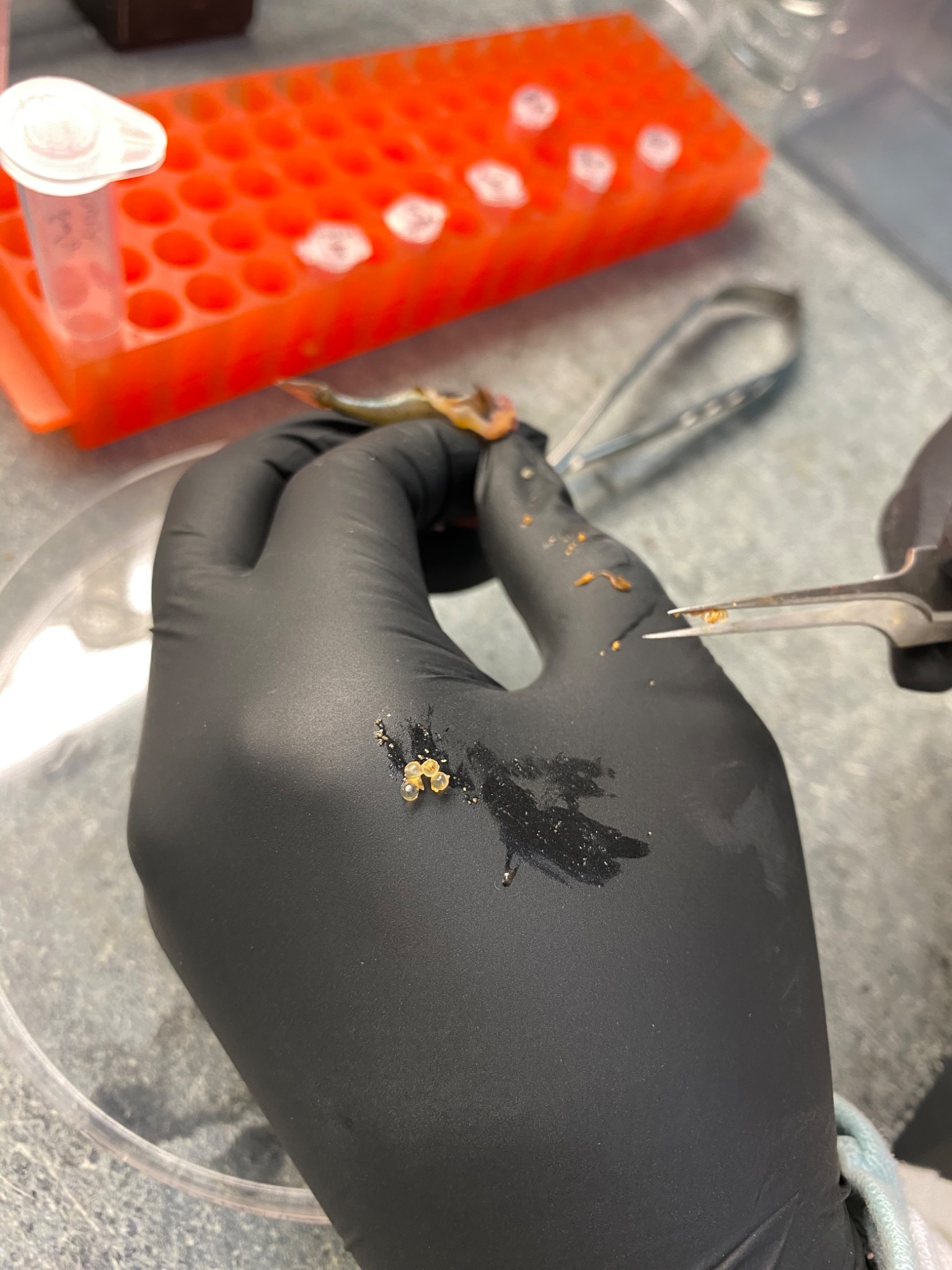 | 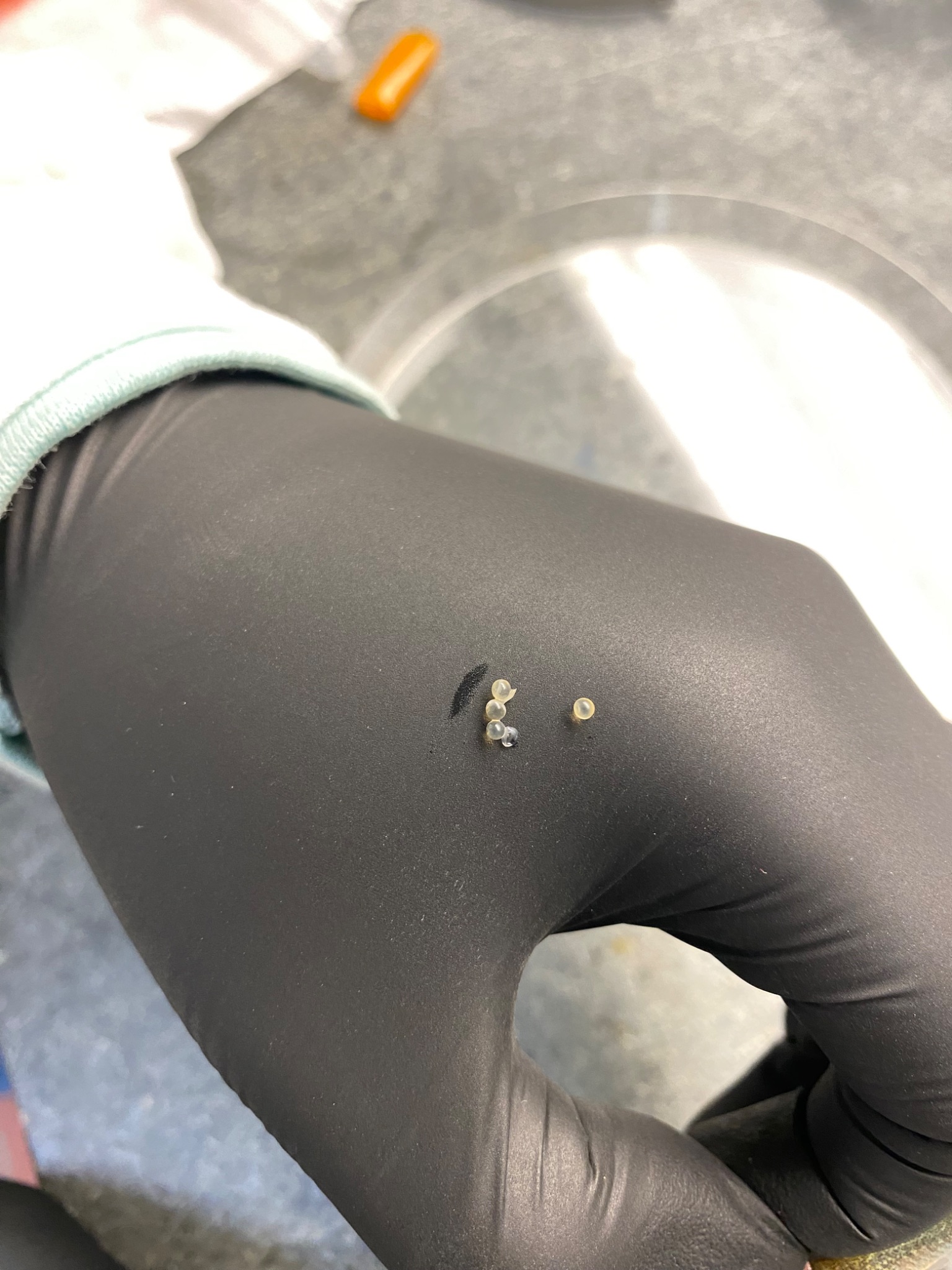 | 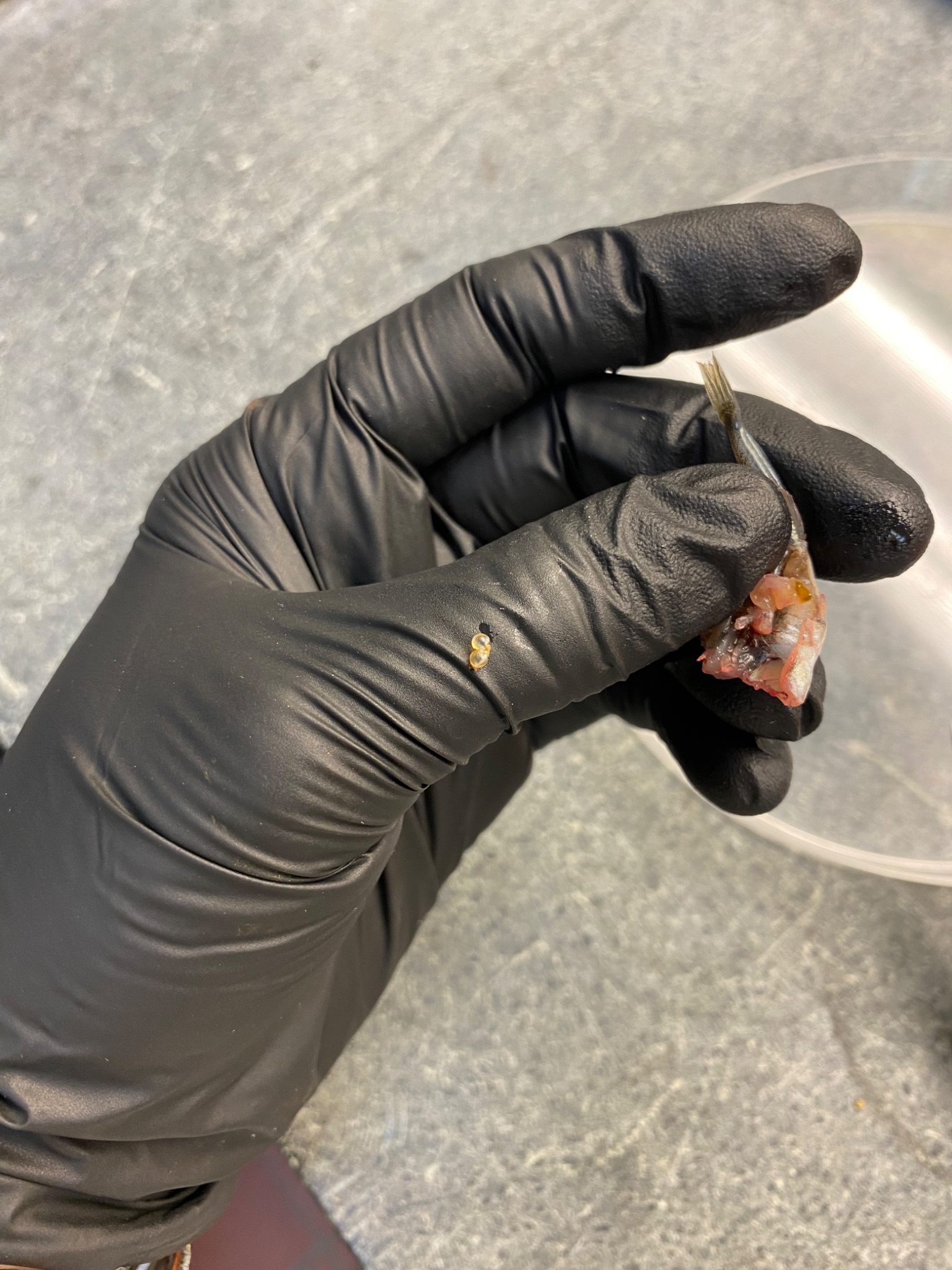 | 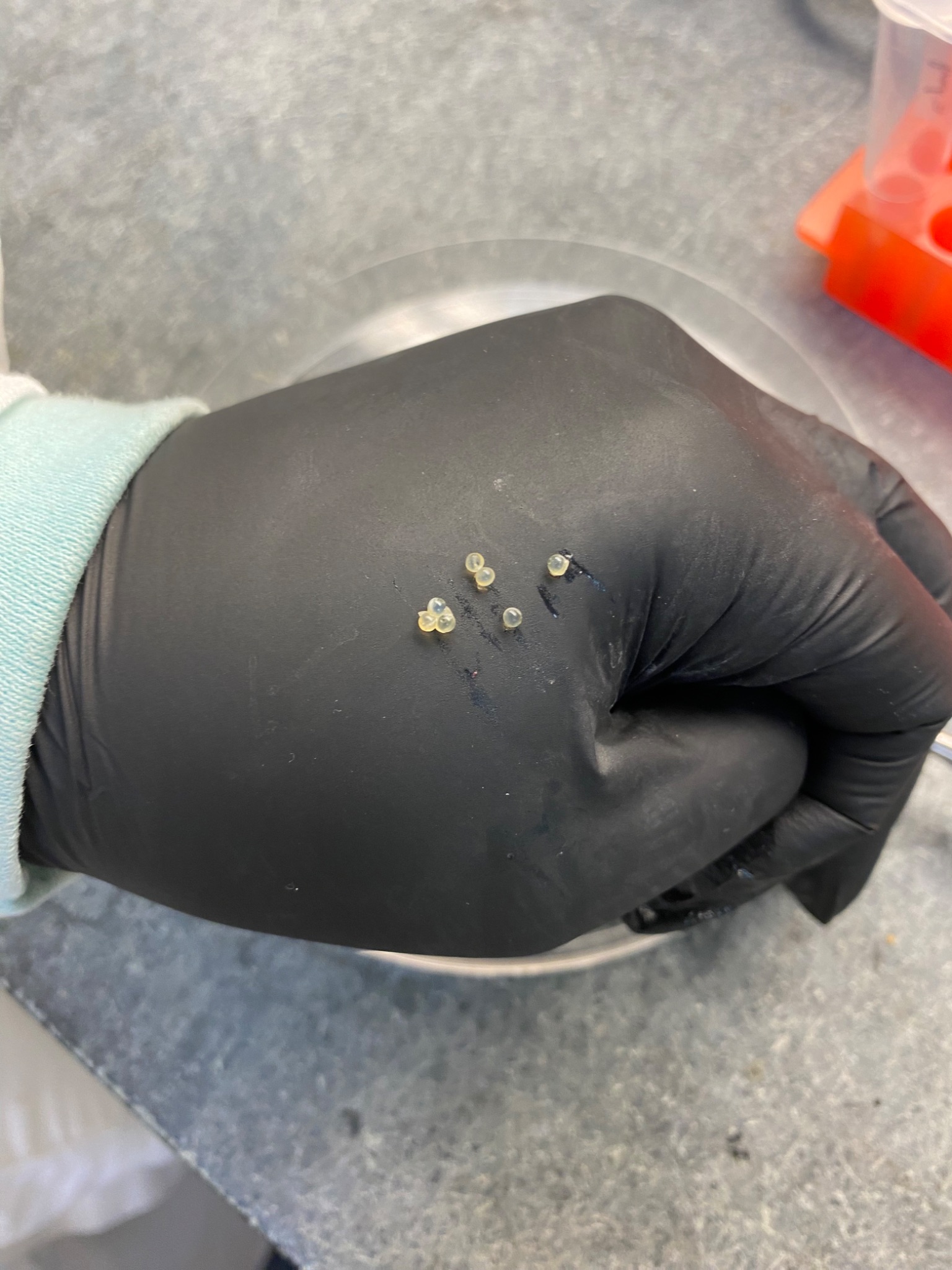 |

**Figure A1. Preliminary data comparing digestion rates of fry and eggs showing fry were digested more rapidly than eggs.** Fry were undetectable in the stomach by one hour post feeding, whereas eggs remained detectable in the stomach at least eight and up to 24 hours.

*Detection of null alleles using Micro-Checker*

Micro-Checker detected evidence for null alleles in our data (table A1), and thus fin clip DNA was amplified and genotyped twice and genotypes that were not the same across runs were excluded from analysis. Combined exclusion probabilities for the panels were calculated using allele frequencies based on screening of parental males in Watson Lake (n = 17 males) and Spirit Lake (n = 12 males). With these relatively low sample sizes, these two populations were not exhaustively sampled, and so the number of alleles as well as their frequencies may be underestimated, and in turn, the exclusion probabilities represent a conservative estimate of the true exclusion probabilities.

**Table A1.** Microsatellite loci used for parentage assignment.

| **Population** | **Locus^1^** | **No. of alleles** | **Null present^2^** | **Null frequency^2^** | **Exclusion probability^3^** | **Combined exclusion probability** |
| --- | --- | --- | --- | --- | --- | --- |
| Watson Lake | Stn45 | 9 | **no** | 0.09 | 0.47 |  |
|  | Stn225 | 12 | **no** | -0.09 | 0.42 |  |
|  | Stn163 | 13 | yes | 0.08 | 0.63 |  |
|  | Stn241 | 9 | **no** | -0.02 | 0.35 |  |
|  | Stn119 | 8 | **no** | -0.06 | 0.36 |  |
|  | Stn344 | 14 | yes | 0.10 | 0.54 |  |
|  | Stn85 | 6 | yes | 0.17 | 0.40 |  |
|  | Stn148 | 6 | **no** | 0.04 | 0.32 |  |
|  | Stn305 | 10 | **no** | 0.03 | 0.48 | 0.995 |
| Spirit Lake | Stn45 | 11 | yes | 0.1515 | 0.52 |  |
|  | Stn225 | 8 | **no** | 0.0309 | 0.33 |  |
|  | Stn163 | 10 | yes | 0.1781 | 0.42 |  |
|  | Stn241 | 7 | yes | 0.254 | 0.32 |  |
|  | Stn119 | 5 | **no** | 0.0871 | 0.21 |  |
|  | Stn344 | 7 | **no** | 0.0792 | 0.42 |  |
|  | Stn85 | 6 | yes | 0.1744 | 0.32 |  |
|  | Stn148 | 6 | **no** | 0.0403 | 0.08 |  |
|  | Stn305 | 6 | yes | 0.1823 | 0.44 | 0.980 |

^1^ See Bay et al. (2017) for primer sequences and microsatellite development

^2^ The presence of null alleles and estimated null allele frequencies were estimated with Micro-Checker software (Van Oosterhout et al. 2004).

^3^ Exclusion probabilities represent the probability of excluding an unrelated individual when one parental genotype is unknown, calculated based on Jamieson & Taylor (1997).
